# Structural Basis of H-NS-Mediated Temperature-Dependent Stimulation of Initial Growth in *Escherichia coli*

**DOI:** 10.64898/2026.03.24.714028

**Authors:** Erika Yamauchi, Yukari Miyake, Kaneyoshi Yamamoto

## Abstract

Bacteria have evolved sophisticated mechanisms to sense and adapt to environmental shifts, particularly when transitioning into the stable, warm temperatures of a host organism. *Escherichia coli*, a primary inhabitant of the warm-blooded host gut, must rapidly initiate growth upon entry to ensure reproductive success. In this study, we demonstrate that the specific growth rate at the onset of the logarithmic phase (defined as μ at *OD*_*ratio*_=0.2) is specifically stimulated across a medium temperature range (30°C–42°C), peaking near the physiological temperature of 37°C. This adaptive response is strain-specific and depends on both the histone-like nucleoid-associated protein (NAP) H-NS and the presence of the Rac prophage. Using high-frequency automated growth monitoring and statistical modeling, we redefine H-NS not merely as a gene silencer but as a critical “nucleoid structural organizer”. Our results indicate that H-NS undergoes a conformational switch at approximately 37°C, transitioning into a parallel form that provides the necessary physical scaffold for nucleoid reorganization. This reorganization is essential for coordinating transcription and replication during the rapid onset of growth. Crucially, we resolve the “silencing paradox”: while H-NS silencing is traditionally thought to be weakened at 37°C, hns-deficient mutants grow significantly slower because they lack this essential structural scaffold. We conclude that the H-NS-mediated physical organization of the genome is more critical for host adaptation than the mere de-repression of the genomic reservoir, enabling *E. coli* to effectively transition into a high-growth state for successful host colonization.

## INTRODUCTION

Bacterial survival depends on the ability to perceive external stimuli and translate them into biological responses. One of the most critical adaptations for *E. coli* is the ability to sense and respond to the temperature of the host gut, which typically resides around 37°C [1]. Success in these environments is often determined by the speed at which a population can initiate growth and expand its clonal presence upon entry. Prophages play a significant role in this adaptive landscape. Approximately 20% of the *E. coli* K-12 genome is composed of prophage DNA, much of which was historically considered “junk” [2,3]. However, contemporary research suggests that these cryptic prophages provide selective advantages, allowing hosts to thrive under various limiting conditions [4]. Specifically, prophages appear to stimulate host growth rates, creating a symbiotic relationship that ensures the reproductive success of both the host and the integrated viral DNA. The modulation of these exogenous genes is primarily governed by H-NS (histone-like nucleoid-associated protein). While H-NS is traditionally characterized as a “silencer” of horizontally transferred DNA [5-8], its role is significantly more complex. Previous evidence suggests it acts as a nucleoid structural organizer required for maintaining a condensed nucleoid structure and facilitating rapid growth [9-12]. The transition from a repressive silencer to an active structural organizer is believed to be temperature-dependent. The objective of this study is to investigate how H-NS mediates the temperature-dependent stimulation of specific growth rate at the onset of the logarithmic phase through structural reorganization. We explore the hypothesis that H-NS-mediated conformational changes, specifically in the presence of prophagic elements like the Rac prophage, provide the physical framework necessary for *E. coli* to adapt to the constant temperatures of warm-blooded hosts.

## MATERIALS AND METHODS

### Bacterial Strains and Genomic Preparation

The bacterial strains used in this study include *E. coli* K-12 MG1655, W3110 type A, and their derivatives (Table 1). W3110 type A was specifically selected as a comparative standard because it carries a complete set of sigma factors and a functional *rpoS* allele, unlike other W3110 laboratory stocks [13].Genomic DNA for W3110 type A was extracted using the Wizard Genomic DNA Purification Kit and whole-genome sequencing was performed via the DNBSEQ-G400 platform to map genomic differences relative to MG1655 [2,3]. The genome of the strain W3110 type A (DRA/DRA009990; RUN/DRR316917) has been published in the DDBJ Sequence Read Archive (https://ddbj.nig.ac.jp//DRASearch/).

### Growth Measurement and Mathematical Definitions

Bacterial growth was monitored using a TVS062CA Compact Rocking Incubator (Advantec) in M9 glucose (M9-Glc) medium at temperatures ranging from 27°C to 45°C. OD_600_ readings were recorded at 15-minute intervals (Fig. 1).

**Figure 1.**
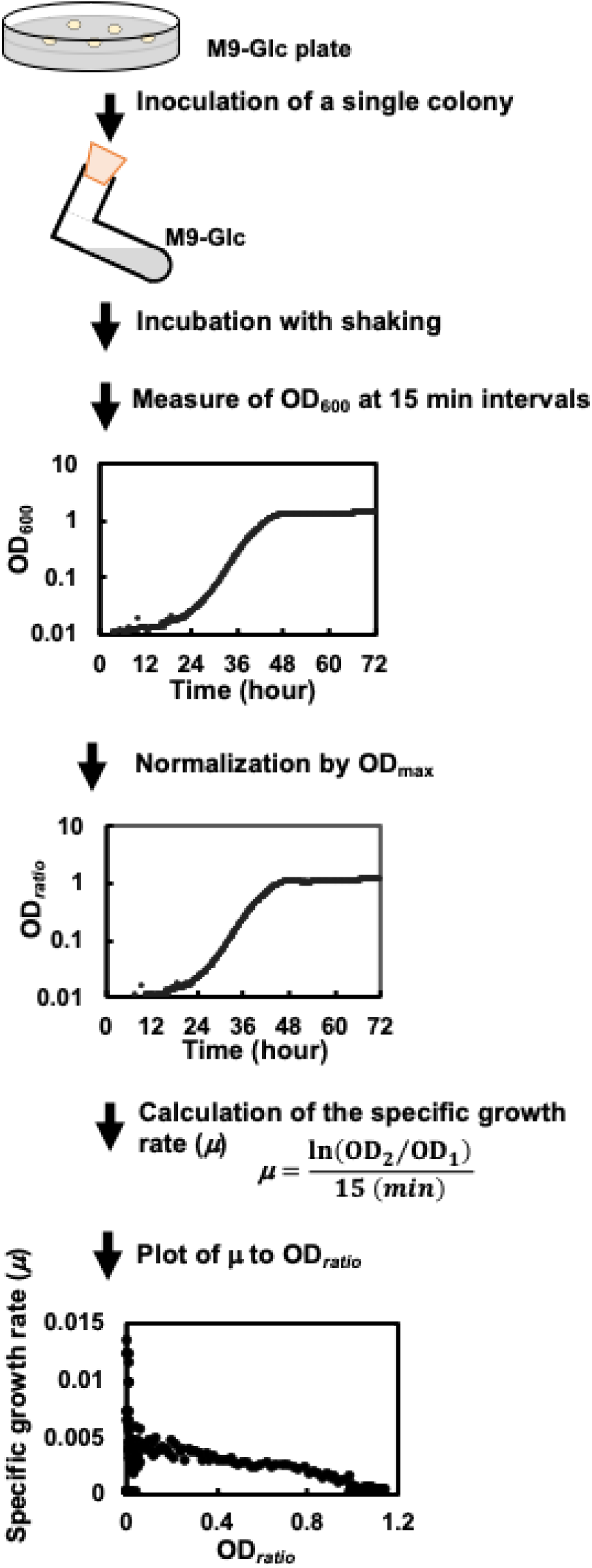
Experimental strategy for evaluating *E. coli* growth kinetics. *E. coli* growth was analyzed by quantifying the relationship between the specific growth rate and culture turbidity. A single colony was inoculated into M9-Glc medium in an L-shaped glass tube and incubated at 37°C with continuous shaking. Growth was monitored via high-frequency OD_600_ measurements at 15-minute intervals for up to 72 hours using a TVS062CA Compact Rocking Incubator (Advantec, Japan). To standardize growth profiles, OD_600_ values were normalized to the maximum observed turbidity (OD_max_) to define the *OD*_*ratio*_ (*OD*_*measured*_*/OD*_*max*_). The specific growth rate (μ) was calculated for each 15-minute interval using the formula μ=ln(*OD*_*2*_*/OD*_*1*_)/15, and subsequently plotted against the *OD*_*ratio*_. Data represent findings from 12 independent colonies per strain.

- **Specific Growth Rate (μ)**: Calculated as *μ = ln(OD*_*2*_*/OD*_*1*_*)/15 min*
- **Normalization**: OD values were normalized to the maximum observed turbidity as *OD*_*ratio*_=*OD*_*measured*_/*OD*_*max*_
- **Specific growth rate at the onset of the logarithmic phase**: Defined as the specific growth rate at the onset of the logarithmic phase. This was calculated as the value of μ when *OD*_*ratio*_=0.2, using a logarithmic fit of the growth curve as *y=aln(x)+b*

### HoSeI Method for Mutant Construction

Mutants for *hns, stpA, hupA, hupB, fis*, and *dps* were created using the Homologous Sequence Integration (HoSeI) method [14]. This technique utilizes CRISPR-Cas9 to induce double-strand breaks at target loci, which are then repaired by integrating a stop codon cassette containing the sequence TAAnTAAnTAA, which terminates all three reading frames. Transformants were selected via antibiotic sensitivity and confirmed through Sanger sequencing (Fig. S2).

## RESULTS

### Temperature-Dependent Growth Patterns

The MG1655 strain exhibited a clear temperature-dependent growth response between 30°C and 42°C (Fig. 2). We observed a maximum specific growth rate at the onset of the logarithmic phase followed by a temperature-dependent rate of deceleration, a phenomenon termed growth compression. At the physiological temperature of 37°C, the specific growth rate and *OD*_*ratio*_ followed the logarithmic fit *y = -0*.*004ln(x)+0*.*0065* (Fig. 2). As the temperature approached 37°C, both the initial velocity and the rate of growth compression increased significantly, indicating a specialized adaptation to host-gut temperatures.

**Figure 2.**
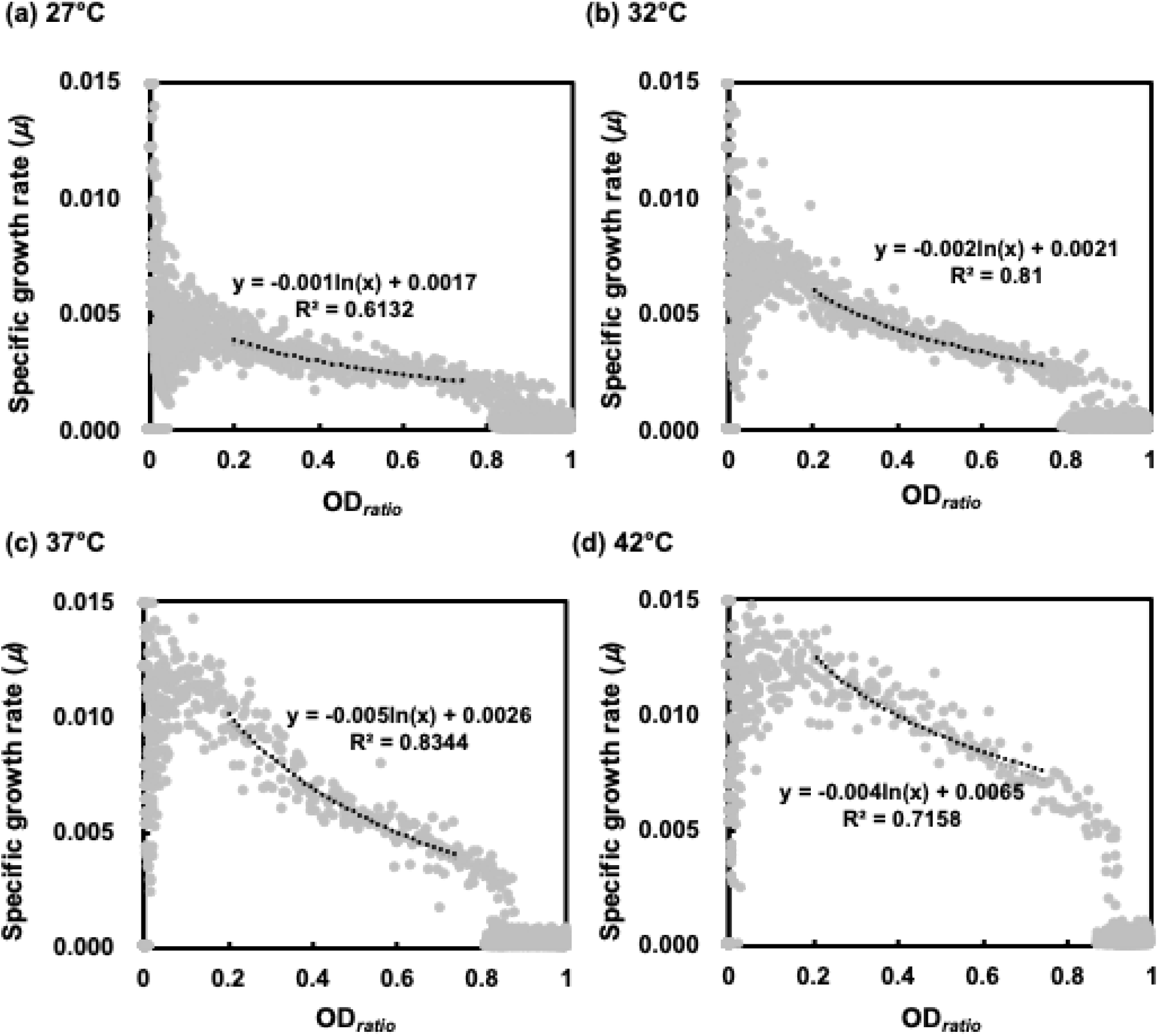
Quantitative analysis of temperature-dependent growth kinetics and the stimulation effect. The relationship between the instantaneous specific growth rate (μ) and normalized culture turbidity (*OD*_*ratio*_) was evaluated across a range of temperatures. (a–d) Representative plots of μ versus *OD*_*ratio*_ for the parent strain MG1655 at 27°C (a), 32°C (b), 37°C (c), and 42°C (d). Individual data points (gray circles) represent measurements taken at 15-minute intervals from 12 independent cultures. The dotted lines indicate the logarithmic regression fit (y=aln(x)+b) calculated within the *OD*_*ratio*_ range of 0.2 to 1.0, which corresponds to the logarithmic growth phase. The regression formula and the coefficient of determination (R^2^) are displayed for each temperature condition. The specific growth rate at the onset of the logarithmic phase is statistically defined as the value of μ at *OD*_*ratio*_=0.2.

### Comparative Analysis: MG1655 vs. W3110 type A

A comparison between MG1655 and W3110 type A revealed distinct kinetic profiles. Despite W3110 type A possessing a functional *rpoS* allele and a complete sigma factor set [13], it failed to match the stimulated specific growth rate at the onset of the logarithmic phase observed in MG1655 at temperatures ≥37°C (Fig. 3a). While MG1655 showed marked stimulation at 37°C compared to 27°C, W3110 type A displayed nearly identical velocities at both temperatures (Fig. 3a). This suggests that the presence of the *rpoS* allele is insufficient to compensate for the lack of specific growth-promoting genomic elements in the W3110 lineage.

**Figure 3.**
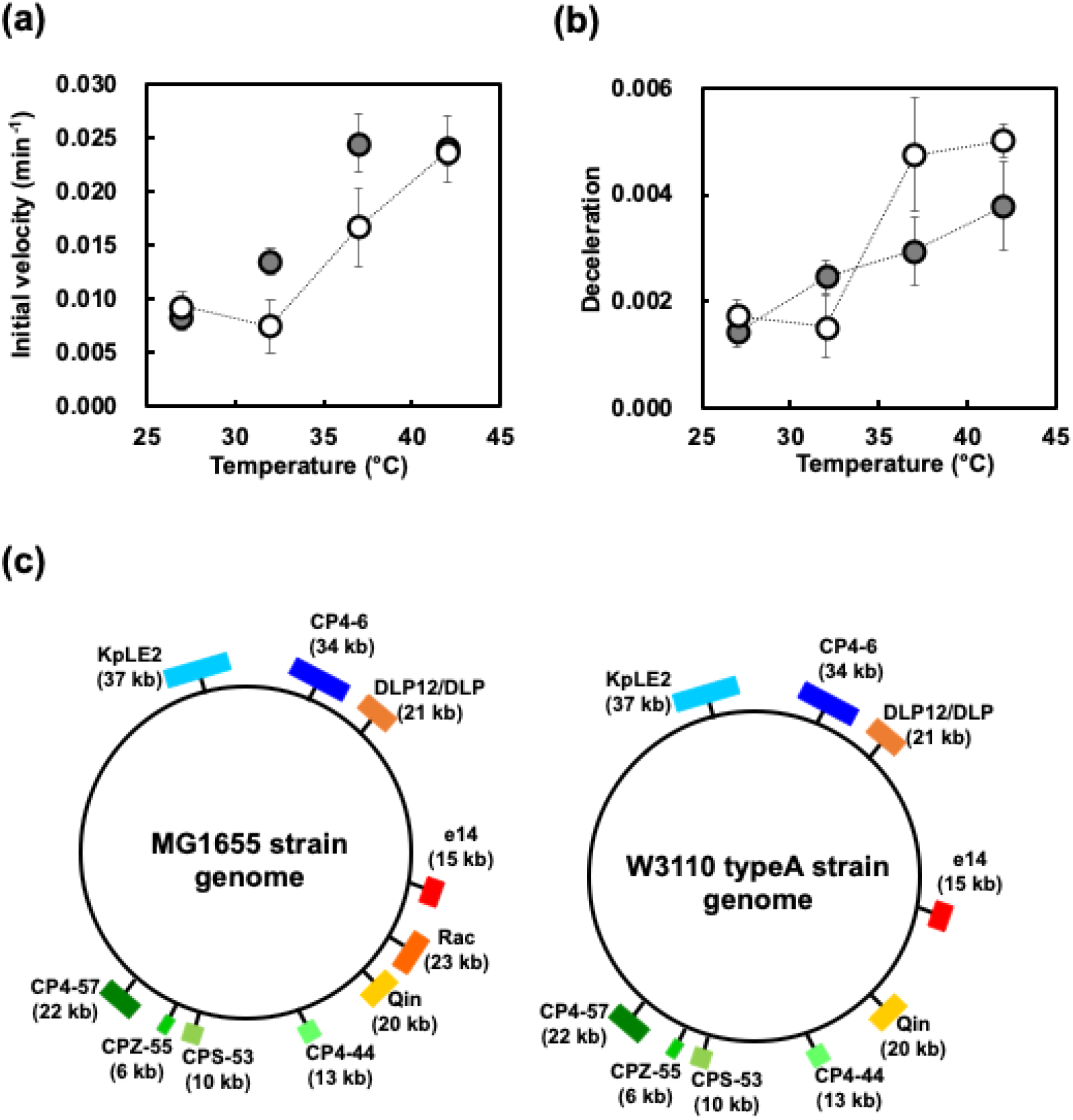
Impact of the Rac prophage on temperature-dependent growth stimulation and comparative genomic mapping. (a) Temperature-dependent profiles of the specific growth rate at the onset of the logarithmic phase (μ at *OD*_*ratio*_=0.2) for *E. coli* MG1655 (filled circles) and W3110 type A (open circles). (b) Comparison of growth deceleration rates across temperatures for MG1655 and W3110 type A. In both (a) and (b), each data point represents the mean value derived from 12 independent colonies, with error bars indicating the standard deviation. (c) Comparative circular genome maps of MG1655 (left) and W3110 type A (right). Colored boxes indicate the positions and sizes (kb) of various prophage elements. The 23-kb Rac prophage (orange box) is present in the MG1655 genome but notably absent in W3110 type A, correlating with the lack of significant growth stimulation at 37°C in the latter strain.

### Genomic Mapping of the Rac Prophage

Whole-genome sequencing of W3110 type A identified a major 23 kb deletion corresponding to the Rac prophage (Fig. 3b). Additionally, a 6 kb region spanning *ynaJ* to *abgA* was found to be absent. Given the differential growth kinetics, these data indicate that the Rac prophage is a primary factor in the stimulation of specific growth rate at the onset of the logarithmic phase at host temperatures.

### H-NS Essentiality and NAP Comparison

To isolate the role of H-NS, we analyzed the initial growth velocities of various NAP mutants. Both MGΔhns and WAΔhns exhibited significantly lower initial growth velocities compared to their parent strains (Fig. 4). In contrast, mutants of *hupA, hupB, fis*, and *dps* did not show a similar essentiality; specifically, *fis* mutants maintained or even exceeded wild-type velocity, and *dps* mutants showed only marginal defects (Fig. 4b). Although *stpA* mutants showed some reduction, the defect in *hns* mutants was the most severe, confirming that the *hns* gene is uniquely required for the temperature-dependent stimulation of specific growth rate at the onset of the logarithmic phase.

**Figure 4.**
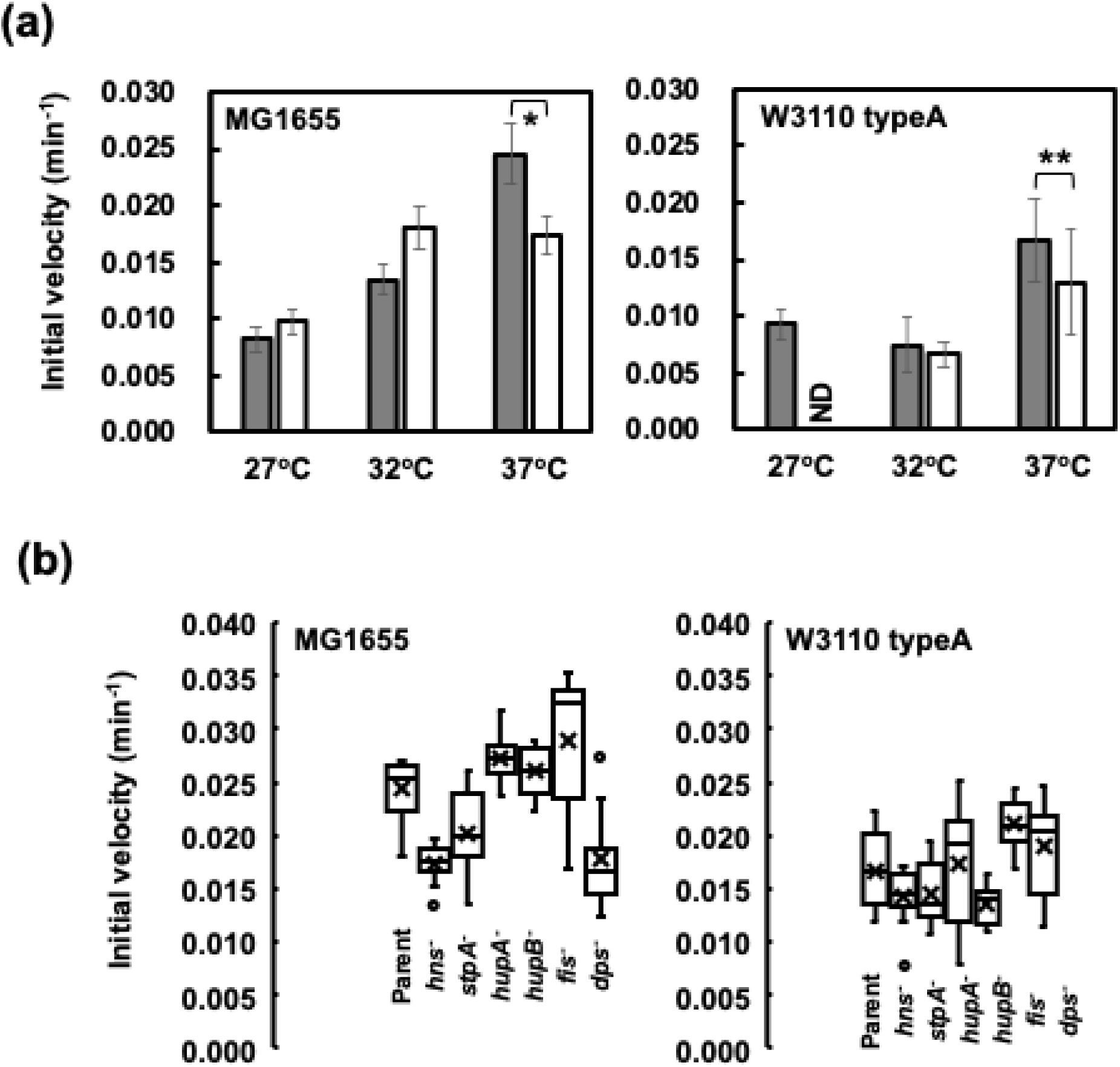
Essential role of H-NS in temperature-dependent growth stimulation and comparison with other nucleoid-associated proteins (NAPs). (a) Comparison of the specific growth rate at the onset of the logarithmic phase (μ at *OD*_*ratio*_=0.2) between parent strains (MG1655 and W3110 type A; open bars) and their respective hns-deficient mutants (MGΔhns and WAΔhns; gray bars) at 27°C, 32°C, and 37°C. (b) Specific growth rates of various NAP mutants (*hns*^*-*^, *stpA*^*-*^, *hupA*^*-*^, *hupB*^*-*^, *fis*^*-*^, and *dps*^*-*^) at 37°C. These results demonstrate that while several NAPs contribute to nucleoid architecture, H-NS is uniquely essential as a nucleoid structural organizer for stimulating initial growth in response to host-gut temperatures. In panel (a), bars represent the mean of 12 independent colonies, with error bars indicating the standard deviation. In panel (b), box plots represent the distribution from 12 independent measurements. Single (*) and double (**) asterisks indicate p<0.05 and p<0.01, respectively, based on Student’s t-test. ND: not determined.

## DISCUSSION

### The Structural Organizer Model

We propose that the stimulation of specific growth rate at the onset of the logarithmic phase is driven by a temperature-sensitive conformational switch in the H-NS protein. H-NS exists in two homodimeric states: an anti-parallel form that induces higher-order multimerization and gene silencing, and a parallel form that promotes unfolding [9-11]. At approximately 37°C, H-NS transitions toward the parallel form, acting as a nucleoid structural organizer [12]. Biochemically, this switch facilitates growth by providing a physical scaffold that coordinates the spatial organization of the nucleoid [12]. This framework is essential for managing replication-transcription conflicts and allowing RNA polymerase access to high-demand regions during the rapid onset of the logarithmic phase. Without this H-NS-mediated scaffold, the Rac prophage cannot exert its growth-promoting effects, even if its genes are de-repressed.

### Addressing the Silencing Paradox

A logical challenge arises regarding H-NS function: if high temperatures weaken H-NS silencing, one might expect *hns* mutants—where silencing is entirely absent—to grow faster at 37°C. However, our data show that *hns* mutants grow significantly slower (Fig. 4). This silencing paradox is resolved by prioritizing the organizer function over the silencer function. While *hns* mutants lack the repression of prophage genes (such as those in Rac), they also lack the essential structural scaffold. Unsilencing a growth-promoting gene is insufficient for stimulating specific growth rate at the onset of the logarithmic phase if the cell cannot physically coordinate the resulting burst in transcription and replication. In the absence of the H-NS organizer, the nucleoid remains uncoordinated, preventing the high-velocity growth seen in the wild type. Thus, the physical organization of the genome is more critical for host adaptation than the mere de-repression of the silenced genomic reservoir.

### Ecological Significance

The H-NS structural adaptation provides *E. coli* with a specialized mechanism to exploit the warm-blooded host environment. By utilizing a temperature-sensitive conformational switch, the bacteria can immediately stimulate their growth velocity upon entering the host gut, ensuring rapid population expansion and successful colonization.

## ACKNOWLEDGEMENTS

We thank Prof. Kenro Oshima of Hosei University for the NGS analysis. We would like to thank Editage (www.editage.com) for English language editing.

## DATA AVAILABILITY STATEMENT

All relevant data are within the paper and its Supporting Information files. The whole-genome sequencing data for *E. coli* W3110 type A have been deposited in the DDBJ Sequence Read Archive (DRA) under accession number DRA009990 (BioProject: PRJDB9626).

